# Sticking around: plant persistence strategies on edaphic islands

**DOI:** 10.1101/2021.06.15.448516

**Authors:** Gianluigi Ottaviani, Francisco E. Méndez-Castro, Luisa Conti, David Zelený, Milan Chytrý, Jiři Doležal, Veronika Jandová, Jan Altman, Jitka Klimešová

## Abstract

1. Species extinction risk at local scales can be partially offset by strategies promoting in-situ persistence. We explored how persistence-related traits of clonal and non-clonal plants in temperate dry grasslands respond intra- and interspecifically to variation in environmental conditions (soil, climate) and insularity.
2. We focused on edaphic island specialist species, hypothesizing that plants experiencing harsh soil environments and strong insularity are distinguished by traits supporting enhanced persistence, such as small stature, long lifespan and resource-conservative strategies. We used linear mixed-effect models and bivariate ordinary least squares linear models to explore the response of species triats to environmental and biogeographic predictors.
3. We found general support for this hypothesis. Soil properties and insularity emerged as the most important drivers of trait patterns. However, clonal species showed more consistent responses to variation in environmental conditions and insularity than non-clonal plants, which were characterized by distinct species-specific responses.
4. Soil properties and insularity confirmed their major role in shaping the persistence strategies of edaphic island plant species. These drivers may exert their effect on specific functions (e.g. belowground resource conservation captured by BDMC). Additionally, we unambiguously identified that clonal species had different persistence strategies than non-clonal ones.

## INTRODUCTION

At the global scale, the rate of species extinction due to environmental changes (e.g. land-use alterations or climate warming) is accelerating at such a pace that the sixth mass extinction has been invoked (Barnosky et al., 2011; Bellard et al., 2012; Ceballos et al., 2015). At finer scales, this process translates into local extirpations of species failing to cope with rapid and abrupt changes (Hylander & Ehrlén, 2013). Plant species may avoid or delay local extinction by surviving outside of their optimal conditions or through various strategies promoting *in-situ* persistence (Marini et al., 2012; Saar et al., 2012). Overall, persistence operates as a counteracting process to local extinction (Auffret et al., 2017). Consequently, analyzing persistence strategies should be critical in conservation biogeography and ecological studies because it may help estimate local extinction risk related to current or anticipated environmental changes.

To successfully persist in an area under changing environmental conditions, plants must be effective in i) acquiring, using and conserving resources (Ackerly & Cornwell, 2007; Saar et al., 2012), ii) occupying new neighboring space through seed dispersal and/or clonal ability (Cody & Overton, 1996; Jiménez-Alfaro et al., 2016), and iii) recovering after disturbance (Martínková et al., 2020; Pausas et al., 2018). How these functions play out under specific circumstances can be assessed through trait-based approaches (Klimešová et al., 2019; Ottaviani et al., 2020a; Pérez-Harguindeguy et al., 2013; Weiher et al., 1999). A good example is offered by seasonal and disturbance-prone temperate grasslands hosting many perennial herbaceous species with different persistence strategies (Klimešová et al. 2016a, 2021; Ottaviani et al., 2020b). A major group comprises long-lived clonal species, capable of both sexual and vegetative reproduction and investing considerable resources belowground into bud-bearing and carbohydrate storage organs (Janovský & Herben, 2020; Klimešová et al., 2016a). Clonal species coexist with many non-clonal species distinguished by shorter lifespan, relying only on regeneration from seeds, and investing fewer resources into bud bank and carbohydrate storage (Martínková et al., 2020). All of these plants can cope effectively with seasonally cold climates and recurrent disturbances (e.g. mowing, grazing, fire; Klimešová et al., 2018, 2021; Ottaviani et al., 2020b). Some of them are also able to deal with increasingly drier seasons, nutrient deposition or altered management regimes (Fischer et al., 2020; Qian et al., 2021). Yet, only those distinguished by high investments into persistence strategies (e.g. bud and seed bank, clonal multiplication) may avoid or delay local extinction caused by ongoing and predicted environmental changes (Marini et al., 2012; Rosbakh & Poschlod, 2021; Saar et al., 2012).

Another potentially critical example in the analyses of persistence strategies is constituted by insular systems. These are “natural laboratories ” (Warren et al., 2015; Whittaker et al., 2017) and suitable models to examine persistence strategies because plants with limited dispersal on steadily isolated islands tend to exhibit adaptive strategies to successfully survive (Cody & Overton, 1996; Conti et al., 2021; Ottaviani et al., 2020a). At the same time, insular systems are particularly vulnerable to species extinctions linked to environmental changes (Courchamp et al., 2014; Macinnis-Ng et al., 2021; Veron et al., 2019). The study of plant functional traits in insular systems has boosted in recent years, providing important insights into the eco-evolutionary dynamics of these systems (e.g. Biddick et al., 2019; Biddick & Burns, 2021; García-Verdugo et al., 2020; Negoita et al., 2016; Schrader et al., 2021; Spasojevic et al., 2014; Taylor et al., 2019). Yet, most of this research targeted dispersal and resource acquisition traits, neglecting other important, non-acquisitive functions (but see Aikio et al., 2020), which are needed for better understanding how plant traits promote persistence and help avoid local extinction (Auffret et al., 2017). For example, plants occurring in highly and steadily isolated places (e.g. distant from species sources preventing or limiting gene flow) should be i) conservative in the way they use resources, ii) grow slower but become older, and iii) allocate more into vegetative reproduction, and when regenerating sexually, producing fewer and heavier seeds with limited dispersal – all strategies indicative of enhanced local persistence (refer to Ottaviani et al., 2020a for an overview).

Trait-based studies are abundant, yet they are often carried out at the interspecific level because differences among species are usually larger than within species (Hulshof & Swenson, 2010; Klimešová et al., 2019; Pérez-Harguindeguy et al., 2013). In the last decade, however, a large body of evidence is showing that intraspecific trait variability can considerably contribute to plant ability to cope with environmental and biotic changes (e.g. Conti et al., 2018; Hulshof & Swenson, 2010; Kichenin et al., 2013; Midolo et al., 2019; Siefert et al., 2015; Violle et al., 2012). Exploring trait patterns at the inter- and intraspecific level may provide a clearer picture of the persistence strategies deployed by insular biota to cope with insularity-related extinction risk – a valuable insight for functional ecology (Weiher et al., 1999) and conservation biogeography (Richardson & Whittaker, 2010).

In this study, we explore links between persistence-related traits at the inter-and intraspecific level with variation in environmental (soil, climate) and biogeographical (insularity) conditions in edaphic islands. These are terrestrial island-like systems defined by the patchy distribution in a landscape of i) discrete bedrock or soil type (e.g. serpentine soils; Harrison et al., 2006; Kazakou et al., 2008; Hulshof & Spasojevic, 2020), or ii) topographic discontinuity (e.g. inselbergs; de Paula et al., 2019; Ottaviani et al., 2016). We focus on edaphic islands constituted by rocky outcrops in Central Europe, which host temperate dry grasslands formed by perennial clonal and non-clonal plant species confined to the outcrops. Due to the nature of any edaphic island system (Kazakou et al., 2008; Spasojevic et al., 2014), we expect soil properties and insularity to play a greater role in shaping persistence-related trait patterns than climate. Because clonal and non-clonal plant species represent distinct life-history strategies in temperate grasslands (Klimešová et al., 2016a; Martínková et al., 2020), we treat them separately. Specifically, we ask:

(Q1) Do clonal and non-clonal plant species consistently exhibit enhanced persistence abilities and more resource-conservative strategies under harsher environmental conditions – i.e. drier and less fertile soils primarily, and, to a lesser extent, warmer and more seasonal climate? If so, do they achieve this through distinct inter- and intraspecific trait responses to the environment?

(Q2) Do clonal and non-clonal plant species experiencing stronger insularity (i.e. growing on smaller and/or more isolated edaphic islands) consistently exhibit enhanced persistence abilities and more resource-conservative strategies? If so, do they achieve this through distinct inter- and intraspecific trait responses to insularity?

## MATERIALS AND METHODS

### Study area and focal species of edaphic islands

We selected 20 rocky (granite or syenite) outcrops occurring in the southern Czech Republic, located at elevations around 450 m a.s.l. (centroid: 49°14 ‘12“N, 15°57 ‘26“E; **Figure 1**). These outcrops (edaphic islands) are scattered across approximately 100 km^2^ and embedded in a landscape matrix of arable land. Their average area is ∼2,912 m^2^ (min = 361 m^2^, max = 14,115 m^2^). The regional macroclimate is temperate (mean annual temperature: 6.5–8°C, annual precipitation: 500–550 mm; Tolasz, 2007), characterized by a marked temperature and precipitation seasonality (Doležal et al., 2021). The vegetation growing on these gently dome-shaped outcrops (maximum elevation ∼5-10 m compared to the surroundings) is acidophilous dry grassland.

**Figure 1.**
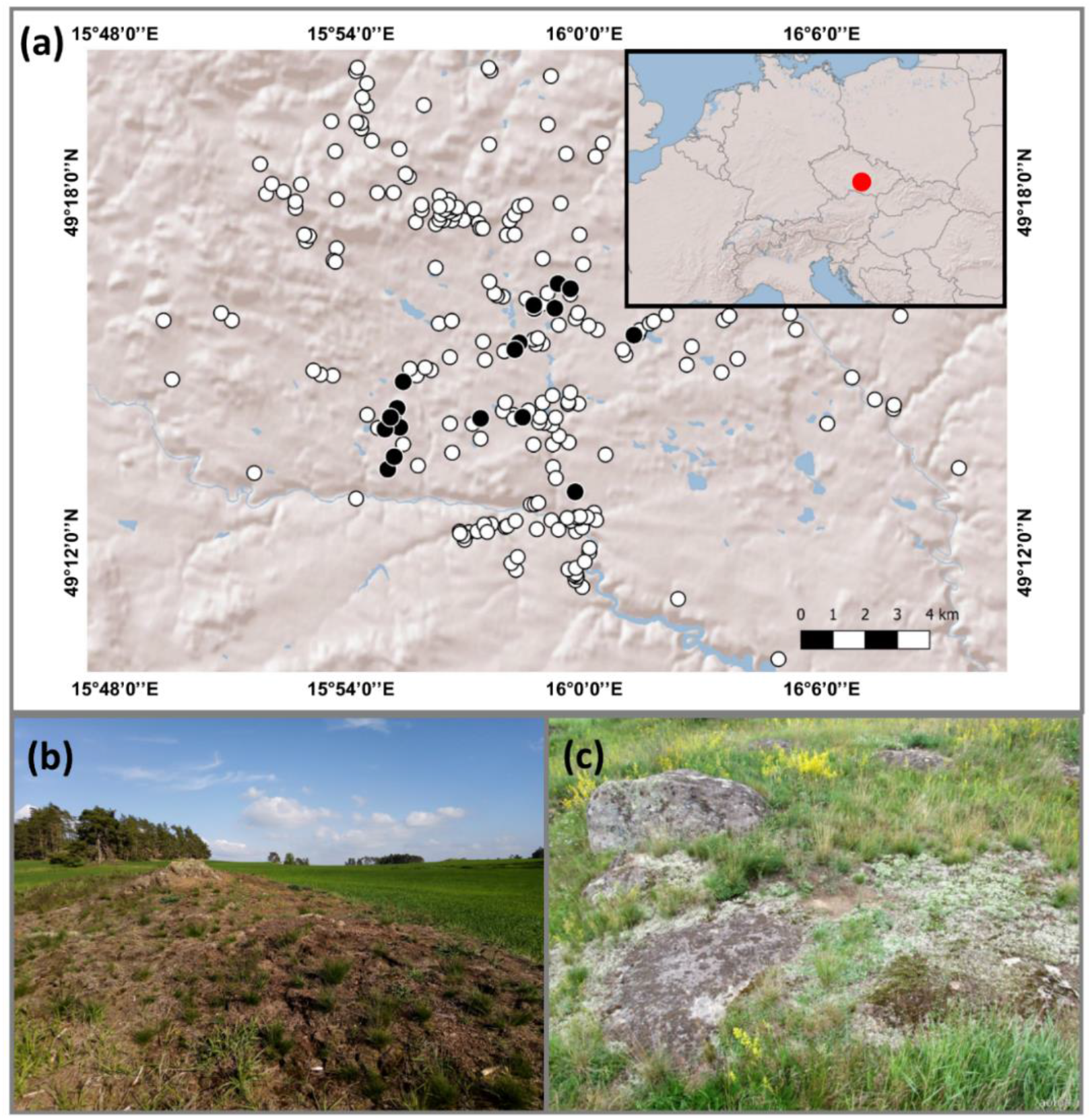
(a) Map and geographical location of the edaphic islands on rocky outcrops in the Czech Republic, Central Europe (red-filled circle in the inset); black-filled circles correspond to the studied edaphic islands, while the white circles represent all the edaphic islands in the surrounding landscape. (b) Example of one edaphic island embedded in the agricultural landscape, and (c) a closer look at the studied shallow-soil, acidophilous temperate dry grassland. Photo credit: FE Méndez-Castro.

We focused on 13 perennial plant species confined to the discrete outcrop grasslands, constituting ∼45% of the total number of specialist species of this vegetation type (Méndez-Castro et al., 2021). These specialist species should be: 1) adapted to the harsh environment of rocky outcrops (especially concerning the putatively strong edaphic filter), and 2) more affected by insularity than non-specialists (for which the surrounding landscape should not be as inhospitable; Conti et al., 2021; Méndez-Castro et al., 2021). The selected species belong to different plant functional types (9 forb species, 2 chamaephytes, 1 grass, 1 sedge), and life histories (5 clonal, 8 non-clonal species). The clonal species are *Carex caryophyllea, Cerastium arvense, Hieracium pilosella, Koeleria macrantha*, and *Trifolium alpestre*, whereas the non-clonal species are *Carlina acaulis, Centaurea stoebe, Helianthemum grandiflorum* subsp. *obscurum, Knautia arvensis, Lychnis viscaria, Scleranthus perennis, Silene nutans*, and *Thymus pulegioides*. Hereafter, we refer to these species using their genus name. Species were found in the majority of the edaphic islands (i.e. 12 to 20 islands), except for *Helianthemum* (6) and *Carex* (9).

### Persistence-related plant traits

During the spring and summer of 2019, we sampled three well-developed and healthy individual plants per species at each edaphic island (total sample size = 538; except in a few instances when only two individuals were available). Trait-data collection and measurement procedures followed standard protocols (Klimešová et al., 2019; Pérez-Harguindeguy et al., 2013). We collected seven persistence-related traits linked to non-acquisitive functions of plants (**Table 1**). In the field, we scored maximum plant height and lateral spread (maximum distance between offspring individuals linked to the parent plant through rhizomes and/or stolons; for clonal species only). For the other five traits, we collected plant material in the field to be processed in the laboratory afterwards. For belowground organ dry matter content (BDMC), a ∼2 cm-long portion of either rhizome (for clonal species) or thick root (for non-clonal species) was cut and put in a paper bag inside a sealed plastic bag. Fresh weight was recorded, then the plant material was oven-dried for 72 hours at 60°C, and the dry weight measured. BDMC was calculated as the ratio between the oven-dry and fresh mass (similarly to leaf dry matter content [LDMC]; Pérez-Harguindeguy et al., 2013).

**Table 1.** Name (indicating if the trait was collected for clonal, non-clonal species, or both), definitions (with variable type, units), relevant functions and key references of the non-acquisitive persistence-related traits included in this study.

| Trait name | Definition<br>(variable type and units) | Function | Key reference |
| --- | --- | --- | --- |
| Plant height<br>[Clonal, Non-clonal] | Maximum plant height<br>(continuous; cm) | Vertical space occupancy;<br>Competitive ability | Pérez-Harguindeguy<br>et al. (2013) |
| Belowground organ<br>dry matter content<br>(BDMC)<br>[Clonal, Non-clonal] | Tissue density of belowground<br>organs, e.g. thick roots,<br>rhizomes<br>(continuous; mg g<sup>-1</sup>) | Resource conservation<br>(water and nutrient) | de Bello et al. (2012) |
| Lateral spread<br>[Clonal] | Distance between offspring<br>individuals and the parental<br>individual in clonal plants<br>(continuous; cm year<sup>-1</sup>) | Horizontal space<br>occupancy; Vegetative<br>reproduction | Klimešová et al. (2019) |
| Age<br>[Non-clonal] | Maximum plant age measured<br>at the root collar<br>(continuous; year) | On-spot persistence | Klimešová et al. (2019) |
| Radial growth<br>[Non-clonal] | Average annual increment<br>measured at the root collar<br>(continuous; µm) | On-spot persistence;<br>Competitive ability | Klimešová et al. (2019) |
| Storage tissue<br>[Non-clonal] | Proportion of tissue for storage<br>measured at the root collar<br>(continuous; %) | Resource conservation;<br>Recovery after damage | Klimešová et al. (2019) |
| Vessel size<br>[Non-clonal] | Cross-sectional maximum<br>vessel diameter measured at the<br>root collar<br>(continuous; µm<sup>2</sup>) | Transport capacity;<br>Growth potential;<br>Resistance to frost and<br>drought | Klimešová et al. (2019) |

Plant anatomical traits – i.e. age, radial growth, storage tissue, vessel size (**Table 1**) – were measured for non-clonal species only (346 individuals analyzed overall). In the field, we cut a ∼2 cm-long fragment located between the root and stem system from each sampled individual. This is indeed the oldest plant part present in the studied growth forms, therefore enables the best estimate of plant age (Klimešová et al., 2019). These fragments were preserved in 50% ethanol, and once in the laboratory, sectioned using a sledge lab–microtome (Gärtner et al., 2015), with thickness between 15 and 40 µm. Cross-sections were then double-stained using a mixture of Astrablue and Safranin dye, dehydrated (with a series of solutions having different ethanol concentrations), washed with xylene and fixed on slides with Canada balsam. These slides were examined at the microscope, and ImageJ software (Schneider et al., 2012) was used to evaluate the number of the annual rings (age), mean annual radial growth, percentage of storage (parenchyma) tissue type, and maximum vessel size in the cross-section.

### Soil, climate and insularity variables

We collected 38 soil samples across 17 edaphic islands (i.e. 2-3 per island depending on its area), close to the location of the climatic data-logger stations (see below). We could not sample all the 20 outcrops due to logistic constraints (see *Data analyses*). Each sample included ∼100 g of soil material. To determine soil nutrient status, we measured eight parameters, namely pH, organic matter, electrical conductivity, total nitrogen, ammonium, nitrate, total phosphorus and exchangeable phosphorus (for details on analytical procedures, refer to **Methods S1**). For soil textural parameters, we measured clay, silt and sand content based on particle size (clay: 0.1-2 µm; silt: 2-50 µm; sand: 50-2000 µm; **Fig. S2**). We also measured soil depth at each sampled plant.

We gathered information on 18 climate variables using 38 automatic climatic stations (TOMST), recording every 15 minutes at 10 cm aboveground and 10 cm belowground for one year (July 2019-July 2020). Like for soil parameters, we located 2 to 3 stations on 17 outcrops. The 18 climate variables relate to temperature and moisture, namely: mean annual value and coefficient of variation (CV) of air temperature, soil temperature and moisture; minimum and maximum value of air temperature, soil temperature and moisture during plant growing season (April to October) and outside the growing season (November to March) (**Fig. S3**).

As a proxy of insularity, we used the target effect metric, which is calculated as the natural logarithm of the ratio between the distance of the target island to its putative species source (i.e. the largest and specialist-richest island) and the square root of its area (Méndez-Castro et al., 2021). Higher values of target effect imply that islands are harder to colonize because of their smaller size or their location far away from the species source (Conti et al., 2021; Méndez-Castro et al., 2021). In the study system, the target effect effectively captures different dimensions of insularity, namely isolation, area, and connectivity (**Fig. S4a**,**b**). The target effect (dimensionless) was calculated for each edaphic island.

### Data analyses

We used the interpolation procedure proposed by Husson & Josse (2016) to impute the missing soil and climate data for three edaphic islands out of 20. Soil properties were aggregated into four predictors: 1) mean depth, 2) depth coefficient of variation (CV), 3) sandiness index, and 4) fertility. Mean soil depth and CV were calculated at the island level using all the measurements taken at each island (i.e. five soil depth measurements/sampled individual = ∼125 measurements/island, total >2500 measurements). The sandiness index was calculated from textural parameters as ∑sand / ∑(silt + clay), i.e. a higher proportion of sand produced higher values of this index, implying less water retention capacity. Soil fertility was identified as the first axis of a Principal Component Analysis (PCA; first axis explaining ∼49% of the variance, with positive scores associated with more fertile soils; **Fig. S5**) ran on the eight soil nutrient parameters. Regarding climate, we conducted a PCA on the 18 variables, and we selected the first two axes to be used as predictors in the models. Specifically, the first PCA axis was positively related to the annual mean of air and soil temperature and negatively associated with the seasonality of those variables (PC1clim; explaining ∼29% of the variance). The second PCA axis was positively related to annual mean soil moisture and less extreme soil temperatures (PC2clim; explaining ∼24% of the variance; **Fig. S3**). For insularity, we included the target effect. Trait values were averaged (using the three values/species/island) for each species at the edaphic island scale.

Before running the models, we controlled for collinearity among the seven soil, climate and insularity variables through Spearman ’s rho correlation coefficient (with Bonferroni correction), and no collinearity issues were detected (**Table S6**). Model-wise, we first ran linear mixed-effect models (LMMs) for clonal and non-clonal species separately. In the LMMs, we set the trait average at the island scale as the response variable, the seven environmental and insularity variables as predictors (i.e. fixed effects; scaled and centered), treating species identity as a random effect (informing on the magnitude of species-specific responses). The variance explained by fixed effects alone (marginal R^2^), and by fixed and random effect together (conditional R^2^) was calculated. Because we aimed at gaining insights into intraspecific responses to different environmental and insularity conditions (and because this study had an inherent explorative component), we examined plant trait-environment links at single trait vs. single predictor level, using bivariate ordinary least squares (OLS) linear models, setting species identity as a grouping factor (i.e. interacting with the single predictor). We identified important relationships based on the model coefficient and its 95% confidence intervals (direction and robustness of the relationship), R^2^ (goodness of fit and strength of the relationship), p-value (significance of the relationship). Except for storage tissue, all the other traits needed log-transformation to accommodate the normality of data distribution and homoscedasticity of model residuals, whereas environmental predictors did not require any transformation. For clonal and non-clonal species separately, we visualized the multivariate trait space identified by functional traits and the species occupancy in this space (i.e. the non-acquisitive persistence niche) through Nonmetric Multidimensional Scaling (NMDS). As traits were measured in different units, we used Gower distance in the NMDS and set 100 random starts and two dimensions. We conducted all the analyses in R version 4.0.1 (R Core Team 2020) using functions available in the packages *missMDA* (for PCA; Husson & Josse, 2016), *vegan* (for NMDS; Oksanen et al., 2017), *lme4* (for LMM; Bates et al., 2015) and *smatr* (for OLS; Warton et al., 2012).

## RESULTS

Predictors alone generally explained a very low proportion of the variability in the LMMs (marginal R^2^ from 1% to 4%), and only for BDMC of clonal species, the fixed effects accounted for 58% (**Table 2**). Most of the model variability was instead explained by the random effect associated with species identity (conditional R^2^ up to 93%; **Table 2**; **Supporting Information 2**). As of NMDS, clonal species (except for *Koeleria* distinguished by consistently taller individuals and *Carex* by higher BDMC) showed a greater overlap in the multifunctional space than non-clonal species, which were characterized by distinct and largely species-specific occupancy of the multifunctional space (**Figure 2**).

**Table 2.**
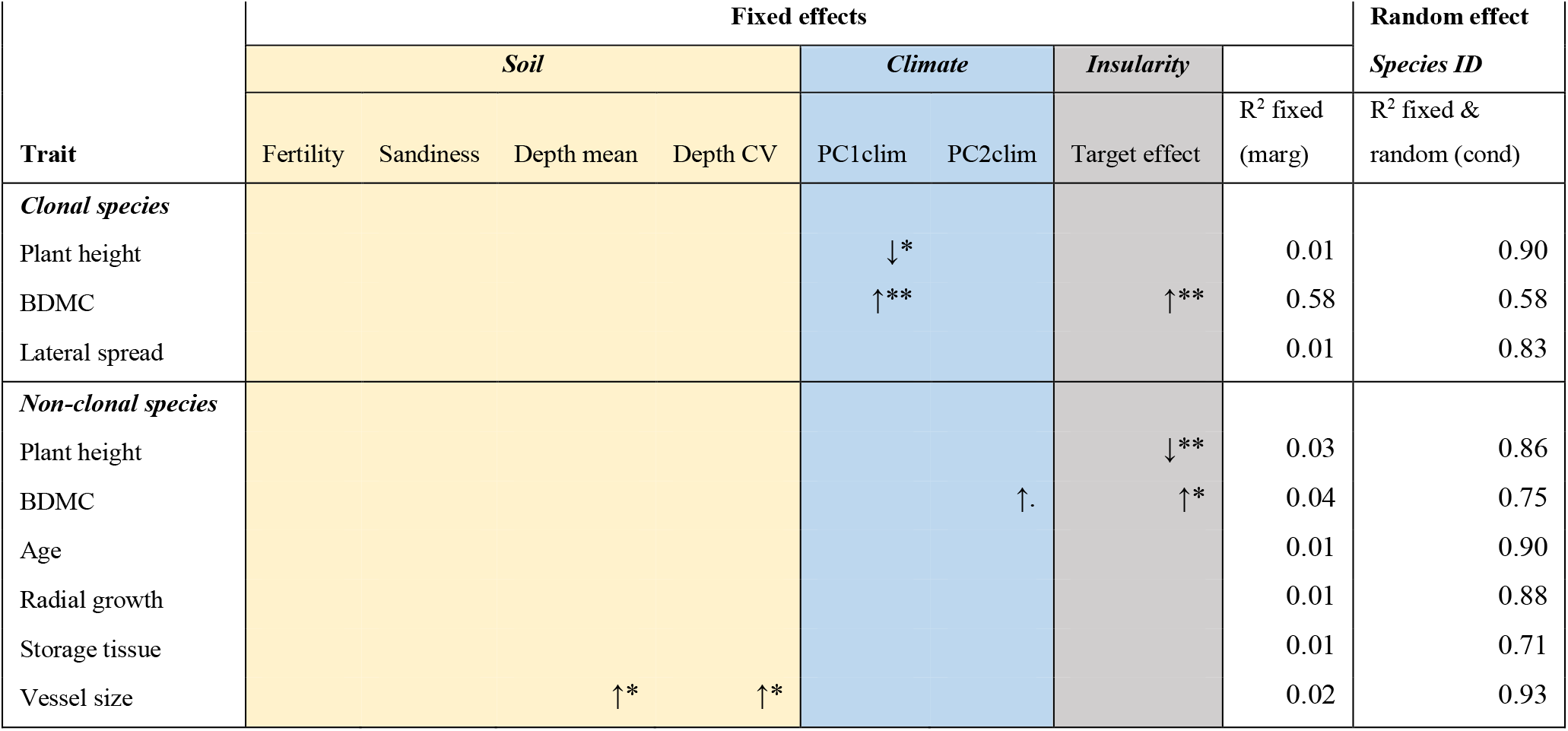
Results of the LMMs for the clonal and non-clonal species separately. Sign (arrow direction), significance (p-value: ** ≤ 0.01; * ≤ 0.05;. ≤ 0.1) and strength (R^2^ values) of single-trait predictor links are reported (marg = marginal R^2^; cond = conditional R^2^). Only the significant relationships are indicated – refer to **Supporting Information 2** for detailed model summary statistics.

**Figure 2.**
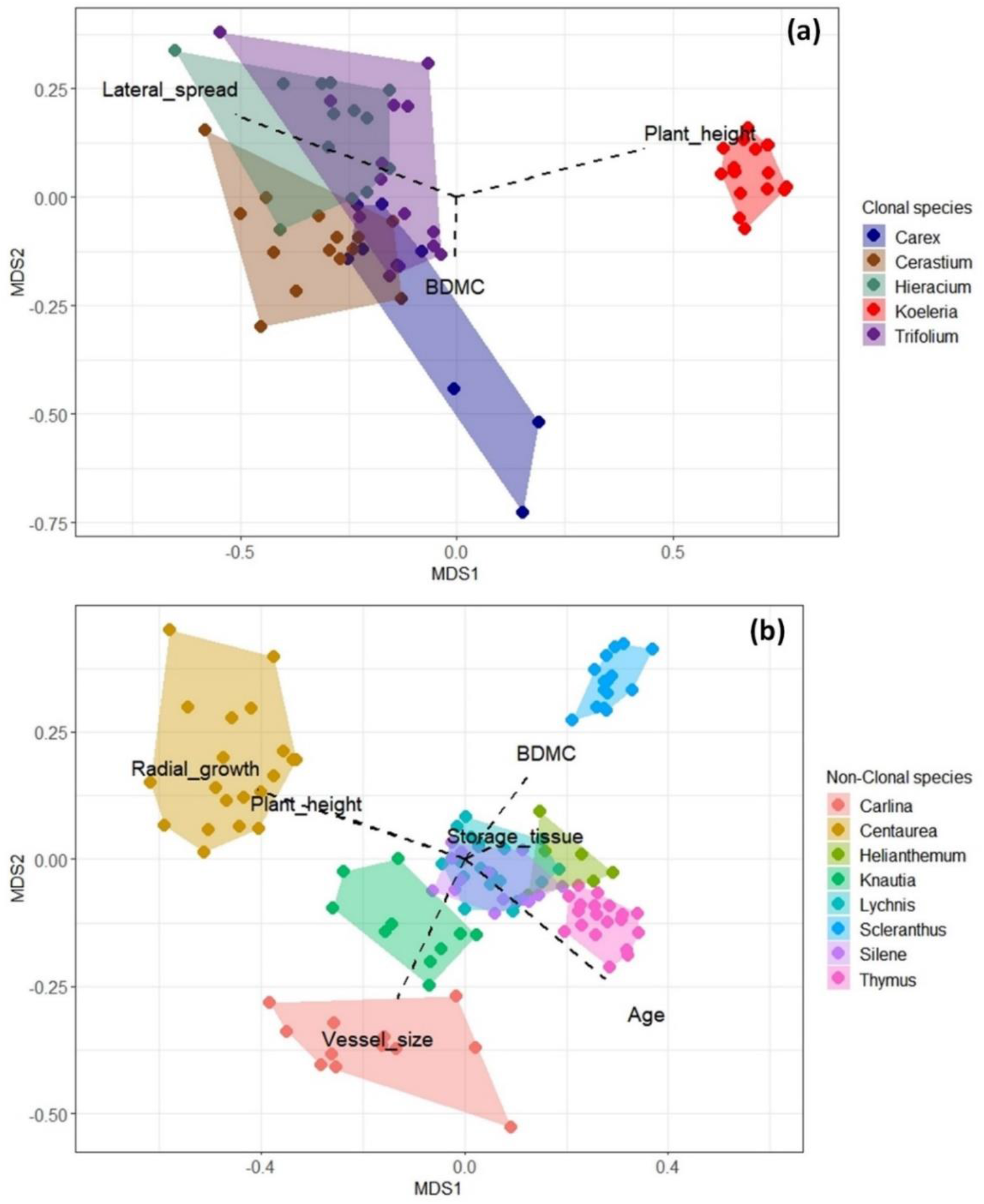
Non-metric multidimensional scaling (NMDS) for clonal (a) and non-clonal species (b), with species occupancy identified by non-acquisitive persistence-related trait values (with direction reported by dashed lines and names). NMDS stress: for clonal species = 0.029, for non-clonal species = 0.127 – both indicating a good fitting of the two-dimensional scaling

Zooming into the links between single traits and single predictors for clonal species, the variance explained by predictors for the most important relationships ranged between 17% and 65%

(**Table 3**). The most consistent relationships were found for the target effect and soil depth CV with BDMC, which exhibited strong positive links across all species. Plant height and lateral spread proved less affected by environmental and insularity factors than BDMC and with less consistent patterns (**Table 3**).

**Table 3.** Results of the OLS regressions for the five clonal species. Sign (arrow direction), significance (p-value: ** ≤ 0.01; * ≤ 0.05;. ≤ 0.1) and strength (R^2^ values) of single trait-single predictor links are reported. Only the most important relationships (based on the model coefficient, 95% confidence interval, significance, R^2^) are indicated – refer to **Supporting Information 2** for further model summary statistics.

| Species | Trait | Soil |  |  |  | Climate |  | Insularity |
| --- | --- | --- | --- | --- | --- | --- | --- | --- |
|  |  | Fertility | Sandiness | Depth mean | Depth CV | PC1clim | PC2clim | Target effect |
| *Carex* | BDMC | ↑** 0.65 | ↓* 0.50 | ↓* 0.61 | ↑* 0.50 |  | ↑* 0.44 | ↑* 0.57 |
| *Cerastium* | BDMC |  |  |  | ↑** 0.44 |  |  | ↑** 0.46 |
| *Hieracium* | Plant height |  |  |  |  | ↓** 0.47 |  |  |
|  | BDMC |  |  |  | ↑* 0.39 | ↑. 0.26 |  | ↑* 0.46 |
|  | Lateral spread |  |  |  | ↑. 0.30 |  |  |  |
| *Koeleria* | Plant height | ↑. 0.17 |  |  |  |  |  |  |
|  | BDMC |  | ↓* 0.26 | ↓. 0.19 | ↑. 0.21 |  |  | ↑* 0.42 |
|  | Lateral spread |  | ↑. 0.20 |  |  | ↑* 0.33 |  |  |
| *Trifolium* | Plant height | ↓. 0.24 |  |  |  |  |  |  |
|  | BDMC | ↑. 0.26 |  | ↓* 0.38 | ↑* 0.38 |  |  | ↑* 0.44 |
|  | Lateral spread |  | ↓. 0.22 |  |  |  |  |  |

For non-clonal species, the most responsive species to changes in environmental and insularity conditions were *Carlina, Helianthemum* and *Lychnis*, whereas *Silene* and *Thymus* responded weakly. The variance explained by the single predictors in the OLSs for the most important relationships ranged between 15% and 87% (**Table 4**). The most responsive traits across species were radial growth and storage tissue. Consistent trait-environment patterns could be identified only across a reduced number of species (i.e. 2 or 3 out of 8); the majority of trait-environment links was highly species-specific, either unique to some species or sometimes contrasting within the same single trait-single predictor relationship (**Table 4**).

**Table 4.** Results of the OLS regressions for the eight non-clonal species. Sign (arrow direction), significance (p-value: ** ≤ 0.01; * ≤ 0.05;. ≤ 0.1) and strength (R^2^ values) of single trait-single predictor links are reported. Only the most important relationships (based on the model coefficient, 95% confidence interval, significance, R^2^) are indicated – refer to **Supporting Information 2** for detailed model summary statistics.

| Species | Trait | Soil |  |  |  | Climate |  | Insularity |
| --- | --- | --- | --- | --- | --- | --- | --- | --- |
|  |  | Fertility | Sandiness | Depth mean | Depth CV | PC1clim | PC2clim | Target effect |
| *Carlina* | Plant height | ↓. 0.27 | ↑. 0.26 |  |  |  |  |  |
|  | BDMC |  |  |  |  |  | ↑* 0.44 |  |
|  | Age | ↑. 0.31 |  |  |  |  |  |  |
|  | Radial growth | ↓. 0.31 | ↑. 0.30 | ↑** 0.57 | ↓. 0.28 |  |  |  |
|  | Storage tissue | ↑* 0.38 |  | ↓. 0.29 | ↑* 0.50 |  |  | ↑* 0.34 |
| *Centaurea* | Storage tissue |  | ↑* 0.27 | ↑. 0.15 |  |  |  |  |
|  | Vessel size |  | ↑* 0.28 |  |  |  |  |  |
| *Helianthemum* | Plant height |  |  |  |  | ↑* 0.71 |  |  |
|  | BDMC |  | ↓* 0.73 |  |  |  |  |  |
|  | Age | ↑* 0.81 |  |  |  |  |  |  |
|  | Radial growth |  | ↑* 0.68 | ↑* 0.71 |  |  |  |  |
|  | Storage tissue |  |  |  |  | ↓. 0.54 |  |  |
|  | Vessel size |  | ↑** 0.87 |  |  |  |  |  |
| *Knautia* | Radial growth | ↓* 0.44 |  |  | ↓* 0.40 |  |  | ↓** 0.61 |
|  | Vessel size |  |  |  |  | ↑* 0.43 |  |  |
| *Lychnis* | Plant height |  |  |  |  |  |  | ↓* 0.22 |
|  | BDMC |  |  |  |  |  |  | ↑* 0.30 |
|  | Age |  |  |  |  |  |  | ↓* 0.25 |
|  | Radial growth |  |  |  | ↑** 0.34 |  |  |  |
|  | Storage tissue |  |  |  |  |  | ↑. 0.16 |  |
|  | Vessel size |  | ↑. 0.16 |  |  | ↑* 0.29 |  |  |
| *Scleranthus* | BDMC | ↑* 0.28 | ↓* 0.30 |  |  |  |  | ↑** 0.47 |
|  | Age |  |  |  |  | ↑* 0.27 |  |  |
|  | Radial growth | ↑. 0.23 |  |  |  |  |  |  |
|  | Storage tissue |  |  |  |  |  |  | ↑. 0.21 |
| *Silene* | Plant height |  |  |  |  |  |  | ↓** 0.46 |
| *Thymus* | Plant height |  |  |  | ↑* 0.25 |  |  |  |

## DISCUSSION

Overall, we found support for our fundamental expectation: soil properties and insularity exerted the most prevalent effects on persistence-related trait patterns. Plants under harsher environmental conditions and in more insular settings were distinguished by resource-conservative strategies. Yet, environmental and insularity predictors generally explained low portions of variability in the models, which was instead mostly related to species identity (**Table 2**). This finding clearly indicates that strategies deployed to persist *in-situ* vary across plant species. This inference was reinforced by the high proportion of variance explained (up to 87%) in the single trait-single predictor models at the intraspecific level (**Table 3, 4**). However, the consistency of trait responses to soil, climate and insularity differed greatly between clonal and non-clonal species. Clonal species tended to show similar trait patterns with overlapping species ‘ non-acquisitive persistence niches, while non-clonal species displayed species-specific trait responses and distinct functional niche occupancies (**Figure 2**).

### Clonal and non-clonal edaphic island plants have different persistence strategies

Except for plant height and BDMC, we assessed different traits for clonal and non-clonal species, mainly because i) collecting anatomical traits for clonal plants is challenging and destructive (the oldest part of the rhizome is necessary but often out-of-reach or decomposed), and ii) some traits may be measured only for one group, such as lateral spread for clonal plants (Klimešová et al., 2019). Yet, all the traits included in this study capture different dimensions associated with non-acquisitive functions shaping species persistence strategies. These are: 1) resource conservation represented by BDMC and storage tissue, 2) plant growth associated with plant height, lateral spread, radial growth, and vessel size, and 3) plant lifespan linked to plant age. Not all traits, however, proved to have the same responsiveness. BDMC emerged as the most responsive and consistent trait across species, especially for clonal plants – the only case where the considerable portion of the explained variability in the model was attributable to predictors only (and no effect of species identity was detected; **Table 2**). This trait is the belowground coarse organ parallel to LDMC and mainly linked to resource conservation function and carbohydrate storage. BDMC can provide insights also into plant ability to overwinter, store water, and multiply clonally – similarly to LDMC reflecting leaf longevity (de Bello et al., 2012; Pérez-Harguindeguy et al., 2013). The functional significance of BDMC and its relationship with other traits would deserve further investigation in different ecological contexts; BDMC may serve as an easy surrogate for other belowground traits and functions that are more laborious to gather data for.

By treating clonal and non-clonal species separately, we were able to detect similarities as well as differences within and among perennial species forming temperate dry grasslands. Clonal species, capable of both vegetative and sexual reproduction, exhibited consistently similar trait responses to variations in environmental conditions and insularity. In temperate grasslands, clonal species tend to prefer moist and nutrient-rich environments (Klimešová et al., 2016b, 2018). Therefore, clonal plants specialized to the dry, sandy, shallow-soil and nutrient-poor environments of isolated rocky outcrops may occupy one of the limits of their ecological niche in the temperate grassland biome. Clonal species may avoid local extinction by extending their longevity through low-cost resource economics (Klimešová et al., 2016a) linked to higher dry matter content of the belowground organ. We cannot, however, fully support this conclusion by direct age determination because for clonal plants, we lack direct measures. Yet, from demographical studies, we know that these species can largely exceed the lifespan of non-clonal plants (Janovský & Herben, 2020), and may persist in remnant populations (Jiménez-Alfaro et al., 2016; Marini et al., 2012; Saar et al., 2012).

A very different scenario emerged for non-clonal species instead. Half of the species (*Carlina, Helianthemum, Lychnis, Scleranthus*) were highly responsive to variation in environmental and insularity conditions. Yet, only in few instances trait responses were consistent across species. Conversely, the other half of the species, especially *Silene* and *Thymus*, remained almost unaffected by marked variations in the edaphic status, climate or insularity. Also, trait responses (or lack thereof) did not mirror plant functional type either. For example, the chamaephytes *Helianthemum* and *Thymus* or the forbs *Carlina, Centaurea, Knautia* and *Scleranthus* were distinguished by well-differentiated strategies to successfully persist on the edaphic islands (**Figure 2b**). This may be due to differences in rooting depth and regenerative strategies (Rosbakh & Poschlod, 2021) and not to plant longevity (see also Doležal et al., 2021). Such a complex and species-specific set of responses may indicate that the focal non-clonal species are likely to be well-adapted to cope with (and not limited by) the distinct ecological and biogeographical conditions provided by the spatially-confined temperate dry grasslands. This inference seems further supported by the almost complete segregation of non-clonal species niches in the multifunctional space identified by non-acquisitive persistence-related traits (**Figure 2**).

### Strong effect of soil and weak climate signal on edaphic island plant persistence

The studied edaphic islands associated with rocky outcrops are distinguished by shallow soils with high sand content, low water retention capacity and low nutrient availability. These harsh edaphic conditions contributed greatly to shaping the non-acquisitive trait patterns influencing the local persistence of perennial plant species. The evidence that soil properties operate as one of the key drivers of persistence strategies edaphic of island plants aligns with other studies (Kazakou et al., 2008; Spasojevic et al., 2014; Hulshof & Spasojevic, 2020). Increasing variability in soil depth was consistently related to higher BDMC values across all clonal species, suggesting that fine-scale soil heterogeneity may push these plants to be more conservative belowground, i.e. a “stay-where-you-are ” adaptive strategy (Graae et al., 2018).

Non-clonal plants grew better in deeper and sandier soils, while plants on more fertile soils tended to grow older but at a slower pace, as for individuals of *Carlina* and *Helianthemum*. These findings challenge the notion that plants inhabiting harsher environments are slower in their growth and longer-lived (Nobis & Schweingruber, 2013). This may suggest that *Carlina* and *Helianthemum* have developed at the intraspecific level unique adaptive strategies to successfully persist in the distinct conditions provided by the edaphic islands, which do not necessarily constrain their growth (Doležal et al., 2021). Non-clonal plants also exhibited a greater ability to transport water through larger vessels in deeper, more variable and sandier soils. Yet, this may be explained by an adaptive hydraulic safety-efficiency functional trade-off (Drake et al., 2015). Larger vessels may imply quicker growth (as found for *Helianthemum*, probably facilitated by its deep rooting ability; Doležal et al., 2021) and higher evapotranspiration, but may also increase the risk of embolism associated with cold and arid spells typical of the highly-seasonal temperate dry grasslands.

Climate affected only a few traits, often showing inconsistent patterns. This flags a minor role of climate in shaping the persistence strategies of edaphic island plants compared to soil properties and insularity. The macroclimate can be considered the same across all the edaphic islands. Yet, microclimate (captured by the data-loggers) can differ within single islands because of variable slope aspect, inclination and solar radiation generated by the rugged terrain and dome-shaped topography of the outcrops (de Paula et al., 2019; Ottaviani et al., 2016). Still, for clonal species, we identified that warmer climates and reduced temperature fluctuations affected plant height negatively and BDMC positively. This result points towards more conservative resource economics under warmer conditions and may indicate that with the anticipated and exacerbating warming of the region, clonal species may be able to cope with these changes (Saar et al., 2012). For non-clonal species, results were even scantier and not pointing towards any generalizable trend.

### Plant trait-insularity links: edaphic islands can operate as true islands, especially for clonal species

Strong insularity promoted belowground resource-conservation strategies across most edaphic island species while tending to reduce plant stature of non-clonal species (i.e. dwarfism; Biddick et al., 2019; Biddick & Burns, 2021; Carlquist, 1974) – both indicative of enhanced persistence abilities (Ottaviani et al., 2020a). Smaller plants with the ability to store resources belowground in dedicated organs (i.e. thick roots, rhizomes) may indicate adaptive strategies towards less costly economics, which may also prolong lifespan (Jiménez-Alfaro et al., 2016; Klimešová et al., 2016a; Saar et al., 2012). These results confirm that insularity can constrain plant growth and promote resource conservation. Additionally, greater insularity affects immigration rates and gene flow (MacArthur & Wilson, 1967; Warren et al., 2015), which may cascade on altered biotic interactions, such as competition, pollination and herbivory (Burns, 2019).

The widespread and consistent evidence that clonal species were more resource-conservative with increasing insularity may provide some insights relevant to their ecology and biogeography. Clonal species are well-known to be poor seed producers (Herben et al., 2015), which may limit their ability to reach distantly-located and/or tiny edaphic islands offering conditions they are specialized for. Yet, once they arrive and establish there, their persistence can be attained via two possible pathways. The first would imply species spreading laterally and multiplying vegetatively and foraging for resources over new areas. The second would involve long-term connection through clonal growth organs, such as rhizomes (Jónsdóttir & Watson, 1997). Both strategies result in an interconnected network of ramets able to share resources among them (Klimešová et al., 2019). The first pathway relates more to the spatial dimension of persistence strategies, while the second refers more to its temporal dimension (Klimešová et al., 2018). The latter pathway seems to be relevant for our edaphic islands, as suggested by high BDMC values (likely linked to a “grow slow-live long ” strategy; Jónsdóttir & Watson, 1997; Klimešová et al., 2016a,b) and no response of lateral spread. Therefore, we may suggest that, especially for clonal species, the studied edaphic islands may be governed by similar ecological and biogeographical forces as true islands (Conti et al., 2021; Méndez-Castro et al., 2021; Ottaviani et al., 2020a).

## Conclusions

In this study, we focused on non-acquisitive traits and functions affecting plant persistence. This topic is largely neglected in functional ecology and biogeography, which typically deal with resource acquisition, dispersal, and sexual reproduction. Our work constitutes one of the first attempts to explore plant persistence-related traits inter- and intraspecifically in insular systems and one of the few to examine changing patterns of these traits across different gradients. Soil properties and insularity confirmed their major role in shaping the persistence strategies of edaphic island plant species. These drivers may exert their effect on specific functions (e.g. belowground resource conservation captured by BDMC). Additionally, we unambiguously identified that clonal species had different persistence strategies than non-clonal ones. The responses of clonal species to major drivers were highly consistent, while non-clonal plants showed distinct and species-specific responses. Lastly, insights into what makes plant species persist *in-situ* can help estimate extinction risk and inform conservation planning of priority elements, such as insular biota.

## ACKNOWLEDGEMENTS

This research was supported by the Czech Science Foundation (project number 19-14394Y to GO, FEMC, LC, VJ; 19-13231S to JK; 19-28491X to MC). GO, FEMC, LC, JD, VJ, JA, JK were supported by the long-term research development project number RVO 67985939 of the Czech Academy of Sciences. We thank Hana Sekerková for her great help with site and species selection and fieldwork campaign.

## AUTHOR CONTRIBUTION

GO conceived the original research idea; GO devised the methodological approach with input from all co-authors; GO, FEMC and LC collected the field samples; VJ conducted laboratory analyses; FEMC measured the insularity metrics; GO and FEMC conducted the statistical analyses; GO and JK led the writing with input from all co-authors.

## DATA AVAILABILITY

The data that supports the findings of this study are available in the supporting information of this article.

## Notes

### Competing Interest Statement

The authors have declared no competing interest.

