## Supporting information 1 for "Sticking around: plant persistence strategies on edaphic islands"

**Article title:** Sticking around: plant persistence strategies on edaphic islands

**The following Supporting Information is available for this article:**

**Methods S1.** *Soil parameters lab-procedures (for nutrient and textural analyses)*

For pH evaluation, a 5 g sample was diluted in 50 ml of distilled water, then shaken by 150 spins/min for 1 hour, centrifuged at 4,000 spins/minute for 10 minutes. This solution was then filtered and pH measured. Electrical conductivity (EC) was measured with the conductivity meter WTW Cond 3110. Soil organic carbon (SOC), total nitrogen (TN) and total phosphorus (TP) were measured using a LiquiTOC tool, while ammonia ($NH_4$), nitrate ($NO_3$) and exchangeable phosphorus (Exch P) using Flow Injection Analysis. For soil textural parameters, we measured clay, silt and sand content based on particle size (clay: 0.1-2 µm; silt: 2-50 µm; sand: 50-2000 µm) using a laser diffraction tool (Fritsch Analysette 22 Microtec plus).

**Fig. S2.** *Ternary plot of soil textural features*

All the 20 edaphic islands fall within three sand-dominated textural categories with low water retention capacity (with high sand content and very low content of clay and silt), namely 1) sand, 2) loamy sand, 3) sandy loam [according to USDA classification].

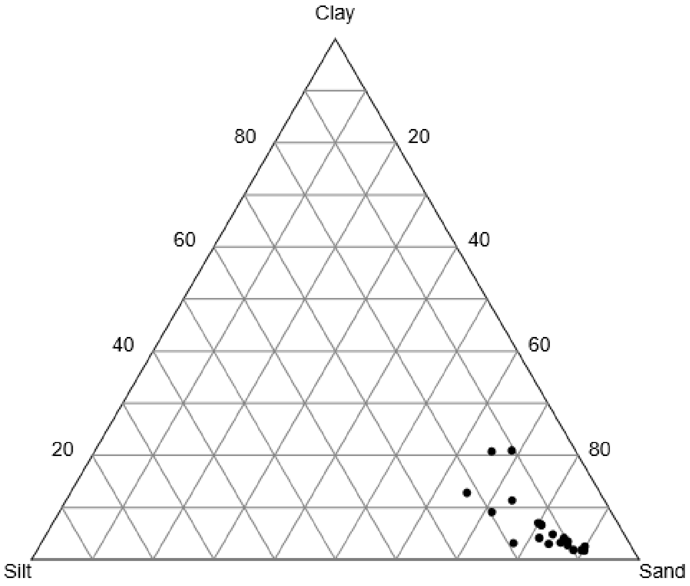

**Fig. S3.** *PCA ran on 18 climate variables*

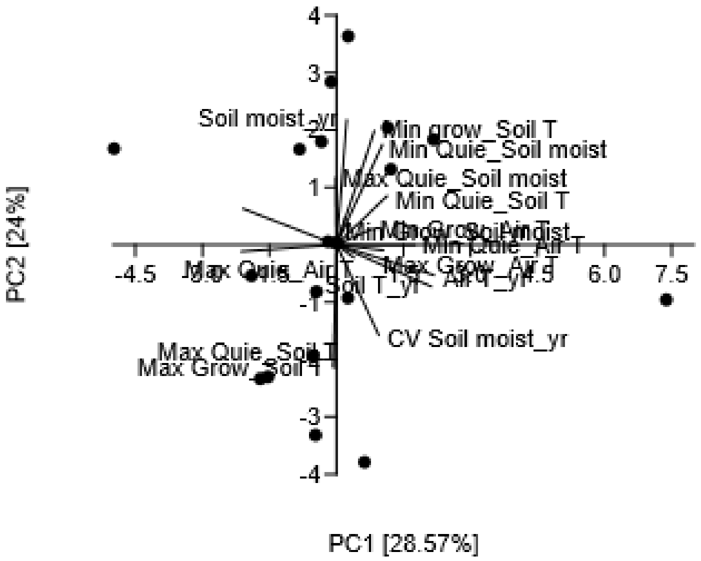


PCA loadings (associated with PC1 and PC2) of the 18 climate variables

| | PC1 | PC2 |
|---|---|---|
| Soil T_yr | 0.41 | -0.14 |
| Air T_yr | 0.42 | -0.10 |
| Soil moist_yr | 0.04 | 0.42 |
| CV Soil T_yr | -0.41 | -0.02 |
| CV Air T_yr | -0.40 | 0.12 |
| CV Soil moist_yr | 0.18 | -0.30 |
| Min Growing_Soil T | 0.16 | 0.38 |
| Max Growing_Soil T | 0.00 | -0.41 |
| Min Growing_Air T | 0.15 | 0.05 |
| Max Growing_Air T | 0.20 | -0.02 |
| Min Growing_Soil moist | 0.04 | 0.09 |
| Max Growing_Soil moist | 0.00 | 0.00 |
| Min Quiescent_Soil T | 0.22 | 0.16 |
| Max Quiescent_Soil T | -0.01 | -0.41 |
| Min Quiescent_Air T | 0.34 | 0.00 |
| Max Quiescent_Air T | 0.07 | -0.04 |
| Min Quiescent_Soil moist | 0.20 | 0.34 |
| Max Quiescent_Soil moist | 0.00 | 0.22 |

**Fig. S4a.** *PCA of the 8 insularity predictors, excluding target effect (from Méndez-Castro et al., 2021) and then the first PCA axis regressed with target effect.*

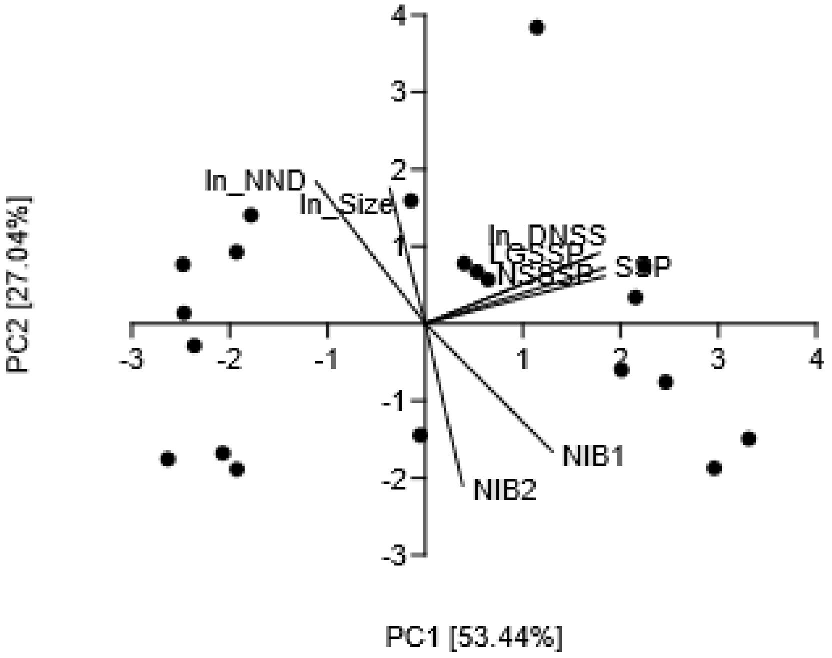


**Acronyms (from Méndez-Castro et al., 2021)**

Size = island size

NND = distance to nearest neighboring island

DNSS = distance to nearest putative species source

NIB1 = number of island in a fine-scale buffer zone (radius 200m)

NIB2 = number of islands in a broad-scale buffer zone (radius 1,000m)

SSP = stepping-stone path

NSSSP = number of stepping stones along the stepping-stone path

LGSSP = largest gap in the stepping-stone path

**Fig. S4b.** Bivariate regression (Standardized Major Axis) between target effect and PC1_scores

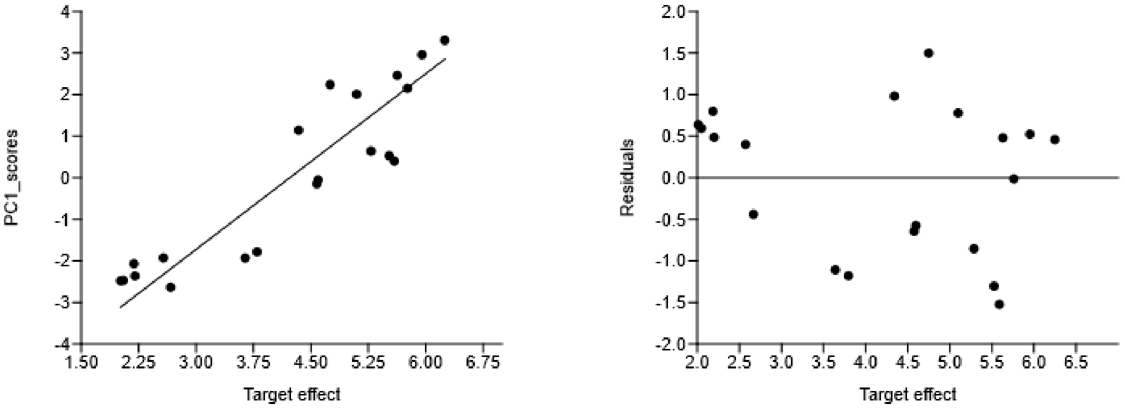


SMA sum stats:

Model coefficient (95% CIs) = 1.41 (1.15, 1.63)

t-value = 10.23; $R^2$ = 0.83; p-value = 0.0001

**Fig. S5.** *PCA ran on 8 soil nutrient parameters (fertility gradient)*

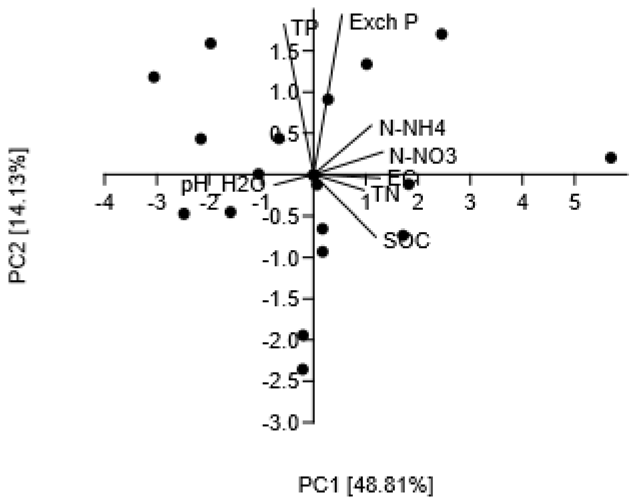


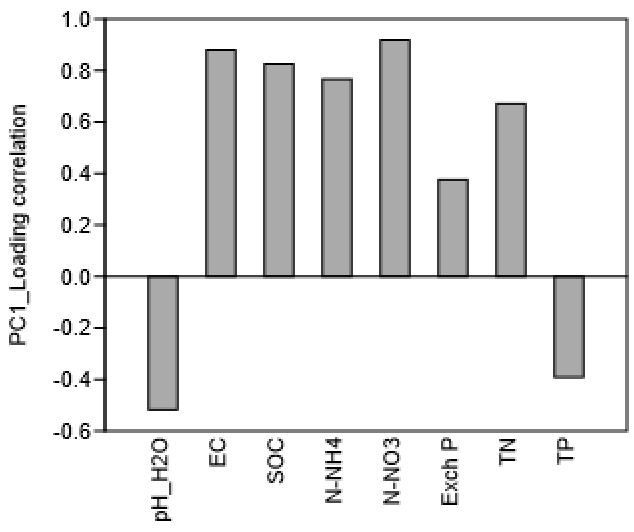

**Table S6.** *Spearman's rho correlation test with Bonferroni correction*

p-values are indicated in the upper diagonal, while correlation coefficients in the lower diagonal. Predictors are all not collinear, i.e. all non-significant correlations, with very low correlation coefficients well below the threshold of 0.75.

| | Soil depth mean | Soil depth CV | Soil sandiness | PC1soil fertility | PC1clim | PC2clim | Target effect |
|---|---|---|---|---|---|---|---|
| **Soil depth mean** | | 1.00 | 0.71 | 1.00 | 1.00 | 1.00 | 1.00 |
| **Soil depth CV** | -0.39 | | 1.00 | 0.08 | 1.00 | 1.00 | 1.00 |
| **Soil sandiness** | 0.48 | 0.29 | | 1.00 | 1.00 | 1.00 | 1.00 |
| **PC1soil fertility** | -0.32 | 0.62 | -0.02 | | 1.00 | 1.00 | 1.00 |
| **PC1clim** | 0.45 | -0.15 | 0.38 | 0.07 | | 1.00 | 1.00 |
| **PC2clim** | -0.12 | -0.23 | -0.25 | -0.14 | 0.25 | | 1.00 |
| **Target effect** | -0.28 | 0.21 | -0.06 | 0.34 | 0.34 | 0.04 | |
